# Fusion transcription factor dosage controls cell state in rhabdomyosarcoma

**DOI:** 10.1101/2025.05.16.654522

**Authors:** Rachel A. Hoffman, Meng Wang, Benjamin D. Sunkel, Thanh Hung Nguyen, Jorge Lopez-Nava, Bishwanath Chatterjee, Wenyue Sun, Frederic G. Barr, Benjamin Z. Stanton

**Author notes:** To whom correspondence can be addressed: (B.Z.S.) or (F.G.B).

## Abstract

In the fusion-positive subset of rhabdomyosarcoma, the PAX3::FOXO1 oncoprotein is the most common fusion driver. We previously established a human myoblast system for inducible expression of PAX3::FOXO1. In the current study, we modulate PAX3::FOXO1 protein expression to understand the epigenetic and phenotypic functions at different PAX3::FOXO1 levels. Proliferative and oncogenic outcomes depend on PAX3::FOXO1 dosage in this system with transformation dominant at intermediate levels and growth suppression dominant at high levels. After prolonged PAX3::FOXO1 expression, there is dosage-dependent heterogeneity in single cell gene expression profiles. We observe a dosage-specific effect for PAX3::FOXO1 chromatin recognition and identify factors that modulate PAX3::FOXO1 chromatin binding. PAX3::FOXO1 dosage affects expression signatures related to cell cycle, epithelial-mesenchymal transition, and myogenesis. Whereas intermediate PAX3::FOXO1 expression maximizes chromatin binding to modulate gene expression, high PAX3::FOXO1 expression alters S phase progression and increases accessibility behind the replication fork. We conclude that PAX3::FOXO1 exerts dosage-dependent functions to influence epigenetic heterogeneity in fusion-positive rhabdomyosarcoma.

## Introduction

To preserve normal function, cells must regulate protein expression levels, starting in the nucleus with chromatin regulatory proteins such as transcription factors (TFs). A subset of TFs called pioneer factors have the ability to bind to motifs within compacted nucleosomal DNA^1,2^ whereas other TFs only recognize accessible DNA.^3^ Key developmental regulators often have pioneer function.^4–10^ Such pioneer factors initially scan chromatin with a transient, non-specific binding mode and then bind nucleosomes through multiple mechanisms.^7,10–13^ Recent studies have revealed that cooperativity between different pioneer factors can fine-tune their recognition of closed chromatin.^10,12,13^ Pioneer factors can also recruit remodeling complexes to increase accessibility.^14^ As suggested by studies of haploinsufficiency, perturbation of pioneer factor dosage in the cell can lead to penetrant phenotypic effects. Dosage sensitivity in a pioneer factor might permit modulation of its recognized sites, with major consequences for cell identity, development, and disease. We hypothesize that genetic and epigenetic changes in human cancers also alter pioneer factor dosage and contribute to disease pathogenesis.

Chromatin replication is a crucial moment in the propagation of the epigenome.^15^ Chromatin must be disassembled ahead of the replication fork and then reassembled behind the fork. Following genome replication, chromatin matures and gains more consistent nucleosome positioning from a more variable initial state in early replication.^16^ Whether pioneer factors that modulate cell identity also modify epigenetic memory during chromatin maturation is currently unknown. Moreover, the effect of varying dosages of a pioneer TF on chromatin replication is not well understood.

Chromosomal translocations resulting in gene fusions that encode chimeric TFs can act as major drivers in cancers. Rhabdomyosarcoma (RMS) is the most common pediatric soft tissue sarcoma and consists of two major subtypes.^17,18^ Fusion-positive (FP) RMS is characterized by *PAX3* or *PAX7* and *FOXO1* fusion events whereas fusion-negative RMS is a heterogeneous category lacking these fusions.^19,20^ PAX3::FOXO1-driven FP-RMS composes approximately 75% of FP-RMS cases and has worse outcomes than PAX7::FOXO1 FP-RMS.^21,22^ No targeted therapies are available for FP-RMS, and standard-of-care treatments have remained the same for decades. FP-RMS is mutationally quiet, with a mutational burden of approximately 0.1 protein-coding mutations per Mb.^23^ To develop new therapies, new insights into this disease are imperative. We previously found that PAX3::FOXO1 has pioneer function, enabling it to bind adjacent to heterochromatin marked with H3K9me3 and induce accessibility.^24^ Moreover, PAX3 is known to bind mitotic chromatin, and PAX3::FOXO1 is highly expressed in the G2 and M phases of the cell cycle.^25,26^ However, whether PAX3::FOXO1 has a role in epigenome duplication and epigenetic memory has yet to be investigated.

Prior studies have investigated the consequences of varying PAX3::FOXO1 expression level. In untransformed murine cell lines, high PAX3::FOXO1 levels induce growth suppression while lower expression favors transformation.^27^ In a FP-RMS mouse model, low PAX3::FOXO1 expression increases tumor-propagating capability.^28^ In human myoblasts expressing PAX3::FOXO1 and MYCN, high expression of PAX3::FOXO1 is necessary for tumorigenesis.^29^ Different levels of PAX3::FOXO1 may direct different oncogenic outcomes, and in human PDX cells, PAX3::FOXO1 mRNA expression varies across single cells.^28^ Understanding the effects of a range of PAX3::FOXO1 dosages is necessary to understand the heterogeneity within and between patient tumors. In this study, we find that PAX3::FOXO1 is a major contributor to the epigenetic state(s) through dosage-regulated chromatin recognition, influencing heterogeneous cell identities and oncogenic character in FP-RMS.

## Results

### Proliferation and focus-forming potential is altered in a dosage-dependent manner by an oncogenic fusion TF in a human myoblast system

For our studies on PAX3::FOXO1 dosage, we used our established system in human myoblasts, Dbt/MYCN/iP3F, that includes MYCN under a constitutive promoter and PAX3::FOXO1 under a doxycycline-inducible promoter (**Figure 1a**).^30^ Myoblasts are myogenic precursors that can develop into mature skeletal muscle cells (myocytes) and share aspects of gene expression signatures in RMS.^17,31–36^ The expression of PAX3::FOXO1 is oncogenic in this system, while its dosage dependencies have remained unclear.^30^ Addition of increasing concentrations of doxycycline for 8 or 24 hours in bulk culture induced increasing PAX3::FOXO1 protein levels (**Figure 1a,b**).^24^ In a two week colony formation assay performed at a range of doxycycline concentrations, we also observe increasing PAX3::FOXO1 levels with increasing doxycycline concentration (**Figure 1b,c**). These data demonstrate that PAX3::FOXO1 expression level can be precisely modulated in Dbt/MYCN/iP3F by the doxycycline concentration in the cell medium and is stable over time.

**Figure 1.**
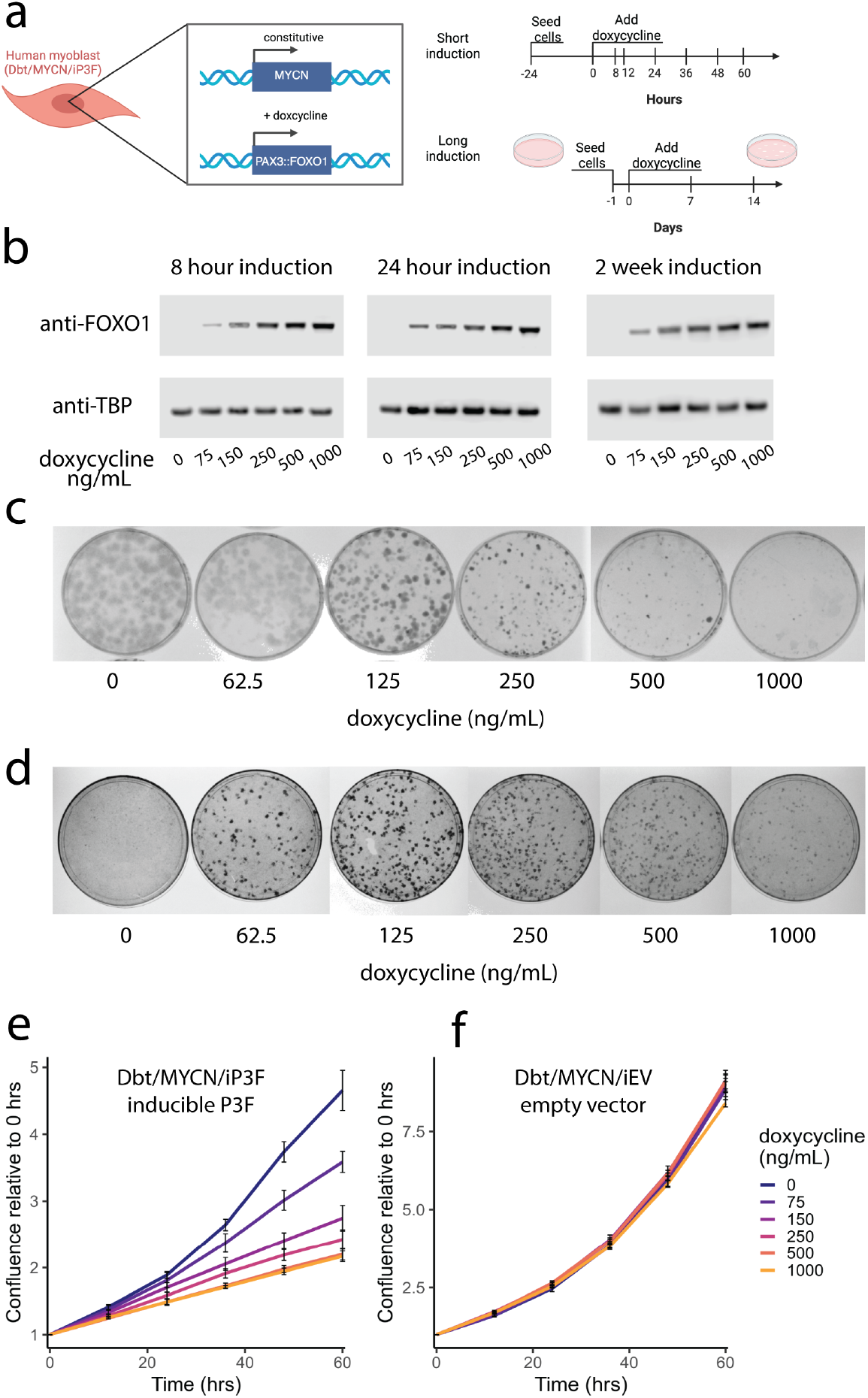
PAX3::FOXO1 dosage alters oncogenicity and proliferation. **(a)** Schematic of our inducible PAX3::FOXO1 system and induction time courses used throughout this study. **(b)** Representative Western blots run on whole cell extracts after a range of doxycycline concentrations were added to Dbt/MYCN/iP3F for 8 hours, 24 hours, or 2 weeks. **(c)** Colony formation assay with Dbt/MYCN/iP3F across a range of doxycycline concentrations. **(d)** Focus formation assay with Dbt/MYCN/iP3F across a range of doxycycline concentrations. **(e)** Dbt/MYCN/iP3F cell confluence measured after 0-1000 ng/ml doxycycline was added to media for 0-60 hours. **(f)** Dbt/MYCN/iEV cell confluence measured after 0-1000 ng/ml doxycycline was added to media for 0-60 hours.

We next asked whether different expression levels of PAX3::FOXO1 result in distinct oncogenic outcomes. To assess this, we performed a focus formation assay across a range of doxycycline concentrations (**Figure 1d**). Maximum focus formation occurs at an intermediate concentration (125 ng/mL). This indicates that increasing PAX3::FOXO1 levels negatively affects oncogenic transformation past a certain point. We also examined the effect of PAX3::FOXO1 dosage on cell proliferation in bulk culture (**Figure 1e**). Any increase in PAX3::FOXO1 expression decreases proliferation compared to the control. This is likely a growth-suppressive effect as observed in prior studies.^27^ Importantly, cell proliferation is not decreased by the presence of doxycycline alone (**Figure 1f**).

We conclude that altering PAX3::FOXO1 expression level modulates cell proliferation and oncogenicity, supporting previous observations.^27,28^ There appears to be an inflection point after which focus forming ability decreases and proliferation is reduced by a consistent level. The timescale of and environment within which PAX3::FOXO1 is expressed also likely impacts these observations. This led us to consider whether there is any difference in dosage-driven gene regulation detectable on a single cell level.

### PAX3::FOXO1 induces heterogenous cell identities reminiscent of human tumors as a function of its dosage

Several single-cell/single-nucleus RNA-seq studies, including a metanalysis of published datasets, have examined RMS patient-derived xenografts and cell lines.^33,34,37–39^ We sought to validate our Dbt model system against these datasets and to investigate single cell gene expression in a PAX3::FOXO1 inducible system. Given that both the time of PAX3::FOXO1 induction and its expression level might be relevant,^24^ we set up experiments varying induction time (8 or 24 hours) while holding dosage constant at 500 ng/mL doxycycline and experiments varying dosage (0, 75, or 500 ng/mL) after a two-week colony formation assay as in **Figure 1c**. We selected 75 and 500 ng/mL doxycycline to induce representative “low” and “high” PAX3::FOXO1 dosage and performed single-cell RNA-sequencing.

We identified and annotated clusters of cells from both the shorter bulk culture experiments and the colony formation assay (**Extended Data Figure 1e-h, see Methods**). In bulk culture at short timepoints (8, 24 hours), there is a strong cycling signature and a ground or baseline state characteristic of G1 phase myoblasts (**Figure 2a, Extended Data Figure 1a**). Strikingly, after a two-week colony formation assay at varying PAX3::FOXO1 dosages, heterogeneity of gene expression is increased and includes cells mirroring states in RMS patient-derived xenografts (“cycling,” “progenitor-like,” “differentiated,” **Figure 2b, Extended Data Figure 1b**,**c**).^34^ Beyond the cell identities seen in prior RMS studies, we also identified cells with high expression of AP-1 TFs and cells with high expression of PAX3::FOXO1 target genes (**Figure 2b, Extended Data Figure 1g**,**h)**.^40^ These results indicate that PAX3::FOXO1 can induce transcriptional heterogeneity in a population of cells over the course of two weeks in a colony formation assay.

**Figure 2.**
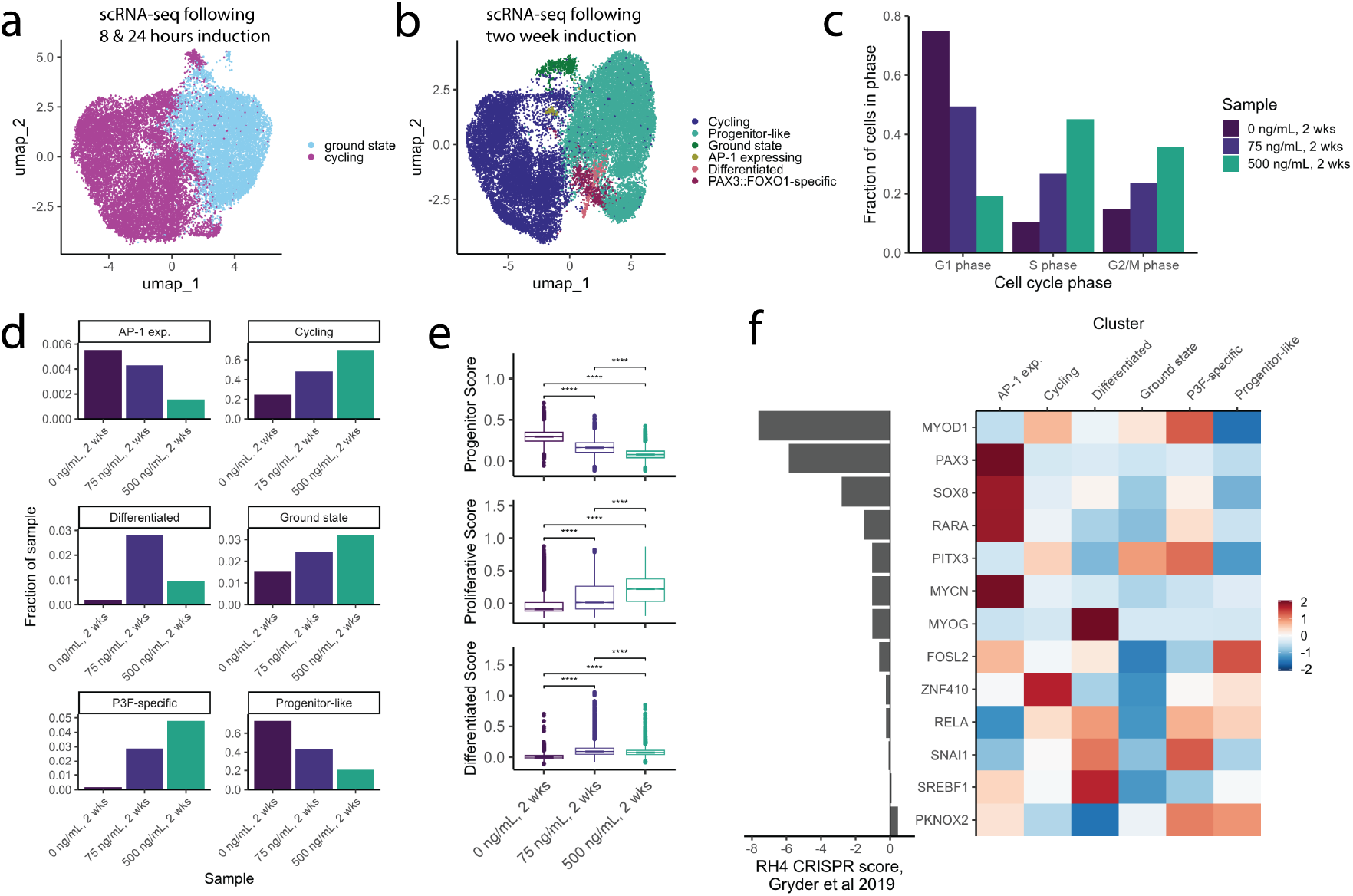
PAX3::FOXO1 expression induces multiple dosage-dependent single cell identities. **(a)** UMAP projection of Dbt/MYCN/iP3F single cell gene expression after induction with 500 ng/ml doxycycline for 0, 8, or 24 hours, with clusters labeled. (**b)** UMAP projection of Dbt/MYCN/iP3F single cell gene expression after 2 weeks of doxycycline induction at 0, 75, or 500 ng/ml, with clusters labeled. **(c)** Fraction of Dbt/MYCN/iP3F cells in each sample in each cell cycle phase based on gene expression. **(d)** Fraction of Dbt/MYCN/iP3F cells in each sample, grouped by 2 week induction cluster. **(e)** Proliferative, progenitor, and differentiated cell identity scores for Dbt/MYCN/iP3F cells in each sample. Cell identity scores were calculated based on Danielli et al 2024. ANOVA with Tukey post-hoc test was used to compare conditions, **** p ≤ 0.0001. (**f)** Right: Aggregate scaled expression of core regulatory TF expression in 2 week clusters. Left: RH4 CRISPR scores from Gryder et al. 2019 indicating essentiality of each core regulatory factor.

We next asked whether altering PAX3::FOXO1’s dosage is instructive for changes in heterogeneity or cell state within our identified clusters. A cell cycle gene expression signature is elevated with increasing PAX3::FOXO1 dosage (**Figure 2c**). High PAX3::FOXO1 expression (500 ng/mL) increases the proportion of S phase and G2/M cells by approximately 25% and 20% respectively compared to cells without induced PAX3::FOXO1 expression. We then examined the distribution of cells in each sample, which reflect a range of PAX3::FOXO1 dosages (**Figure 2d, Extended Data Figure 1d**). Samples with PAX3::FOXO1 induction have more cells in the “PAX3::FOXO1-specific” cluster, agreeing with expectations. In patient-derived scRNA-seq, “progenitor-like” identity is more enriched in FN-RMS tumors and differentiated (myogenic) identity is associated with FP-RMS.^34^ Similarly, we observe only 20% of cells in the high PAX3::FOXO1 induction sample with progenitor-like identity, while approximately 60% of cells in the uninduced sample are in this cluster (**Figure 2d**). Interestingly, the uninduced sample has the highest proportion of “AP-1 expressing” cells (0.5%). “Differentiated” cell identity makes up approximately 3% of the low PAX3::FOXO1 sample, compared to <1% of the other samples. Our results indicate that PAX3::FOXO1 expression induces cellular heterogeneity in a dosage-dependent manner, with similarity in our cell populations to populations found in patient samples.^34^

Next, we calculated RMS cell identity scores identified by Danielli et al in each cell in our dataset and analyzed these scores with respect to PAX3::FOXO1 dosage (**Figure 2e**).^34^ As dosage increases, the progenitor score decreases while the proliferative score increases. Similarly, the differentiated score is low in the absence of PAX3::FOXO1 induction, highest at low PAX3::FOXO1 expression (75 ng/mL), and decreases again at high PAX3::FOXO1 dosage. We note that “differentiated” identity is associated with drug-resistant persister cells.^28,33,34^ To drive these different gene expression patterns, we hypothesize that PAX3::FOXO1’s chromatin recognition may differ across varying PAX3::FOXO1 expression levels.

A set of core regulatory transcription factors (CRTFs) have been identified as part of the core regulatory circuitry (CRC) that establishes epigenetic cell state in FP-RMS, and we wondered how they might underlie the observed single cell identities.^41^ We investigated whether expression of the CRTFs varies across single cells after two weeks of PAX3::FOXO1 induction.^41–43^ We found that their expression is not uniform across the single cell clusters and is instead highly variable (**Figure 2f**). For example, SOX8 is most highly expressed in the “AP-1 expressing” cluster. Some CRTFs are highly expressed in multiple clusters (MYOD1, PITX3, RELA). Interestingly, the essentiality of the CRTFs for proliferation in RH4 cells (“RH4 CRISPR score”, ref. 40) does not appear tied to high expression across the majority of cell populations as might be expected. We suggest that in FP-RMS, the expression of CRTFs may vary within a heterogenous population of single cells as opposed to one unifying CRC across all cells. Ultimately, the observed heterogeneity raises the question of how these cell identities and their distributions are established epigenetically by PAX3::FOXO1.

### PAX3::FOXO1 chromatin binding is dosage-sensitive in human myoblasts

To investigate how PAX3::FOXO1’s chromatin recognition might mediate the observed dosage-dependent effects, we induced PAX3::FOXO1 expression with multiple doxycycline concentrations during a two week colony formation assay. We then used our established approach to assess PAX3::FOXO1 genome binding through FOXO1 chromatin immunoprecipitation.^24^ We observed that some PAX3::FOXO1 sites are occupied across all dosages. For example, *FGFR4* is a key regulatory target of PAX3::FOXO1 and has two PAX3::FOXO1 regulatory sites downstream of the gene body across dosages (**Figure 3a**).^40^ However, the total number of PAX3::FOXO1 ChIP-seq peaks, or significant genomic binding sites, at each dosage varies widely (**Figure 3b**). There are only 1,261 consensus peaks, or reproducible peaks across independent experiments, identified across replicates at 0 ng/mL. It is technically possible that this may indicate low PAX3::FOXO1 expression in the absence of doxycycline that is undetectable by immunoblotting. However, once doxycycline is added the number of PAX3::FOXO1 sites increases to 24,240 sites at 75 ng/mL. The maximum number of consensus PAX3::FOXO1 sites (39,703) occurs at 150 ng/mL. The number of consensus PAX3::FOXO1 sites then decreases with increasing dosage. PAX3::FOXO1’s binding specificity may be modulated based on the average amount of PAX3::FOXO1 protein within each cell.

**Figure 3.**
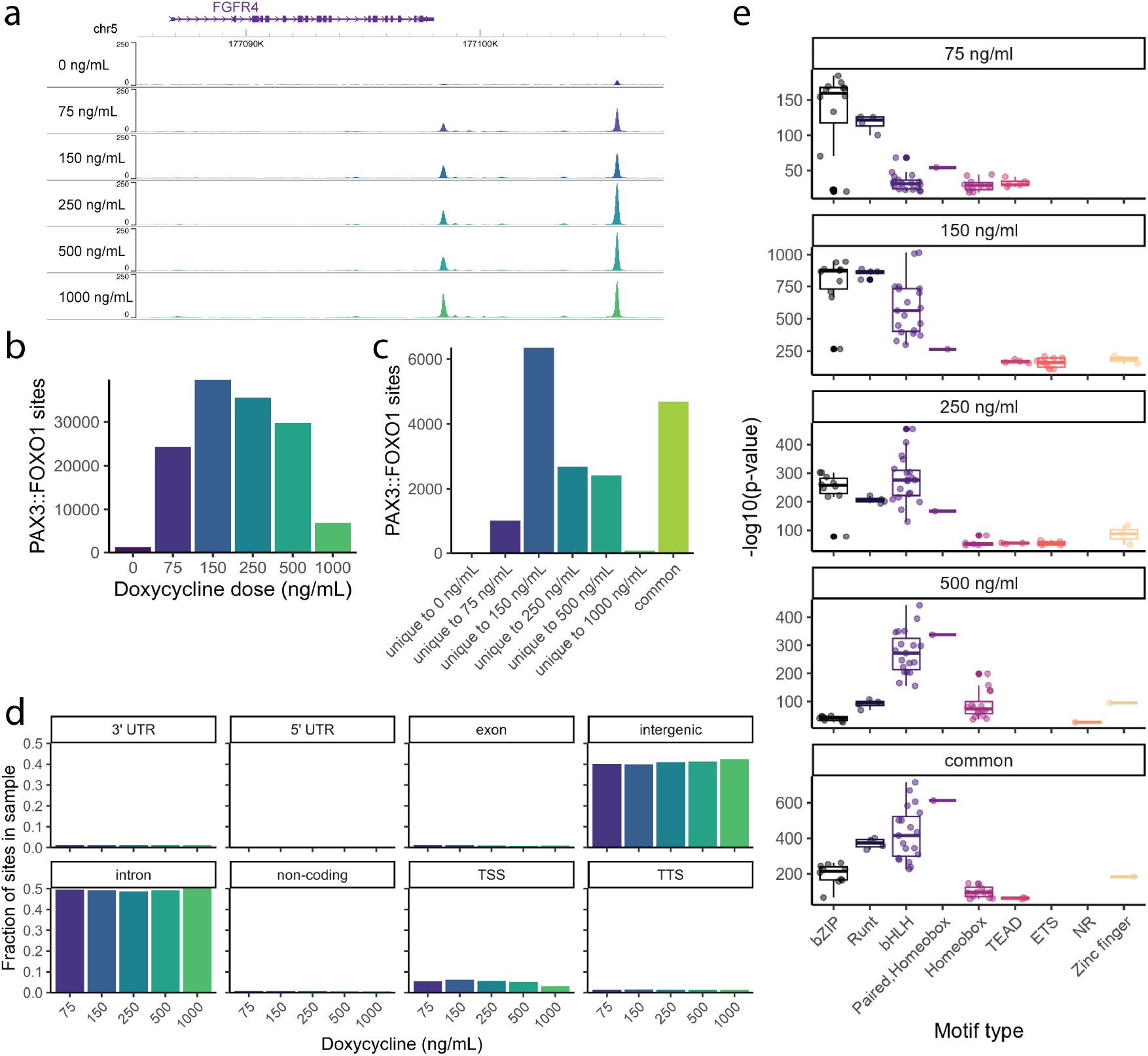
PAX3::FOXO1 dosage modulates its chromatin binding. **(a)** anti-FOXO1 (PAX3::FOXO1) ChIP-seq signal at *FGFR4* in Dbt/MYCN/iP3F with doxycycline induction at 0-1000 ng/ml for 2 weeks. **(b)** Number of consensus PAX3::FOXO1 ChIP-seq peaks at each doxycycline concentration. **(c)** Number of unique PAX3::FOXO1 peaks to each condition and number of common peaks across all induced conditions. **(d)** Fraction of PAX3::FOXO1 peaks at each doxycycline concentration in each genome annotation category. **(e)** Motif enrichment within unique PAX3::FOXO1 peaks to each doxycycline concentration and within common peaks across all doxycycline concentrations. Each motif found is plotted by the -log10 p-value and motif family determined by HOMER.

To gain insight into dosage-specific chromatin recognition mechanisms, we categorized all PAX3::FOXO1 peaks as unique to one dosage (“dosage-specific,” i.e. peaks that are significant at only one dosage), occurring in all dosages (“common,” i.e. peaks significant across all dosages), or occupied in a subset of dosages. We took a conservative approach, identifying sites as occupied in a given dosage by the presence of a significant consensus PAX3::FOXO1 ChIP-seq peak. The maximum number of dosage-specific sites, 6,361, occurs at the intermediate doxycycline level 150 ng/mL (**Figure 3c**). There are also 4,688 sites occupied across all the induced (75-1000 ng/mL) conditions (“common” in **Figure 3c**). At PAX3::FOXO1 sites occupied across either two or three dosages, the trends are similar (**Extended Data Figure 2a**,**b**). To probe distinctions in the identified PAX3::FOXO1 sites across dosages, we assessed the genomic feature annotations of PAX3::FOXO1 sites and found that there are no substantial changes in distribution across categories of genomic features based on dosage (**Figure 3d**). In summary, PAX3::FOXO1 genomic binding is maximized at intermediate dosages, suggesting there are differences in PAX3::FOXO1 chromatin recognition based on PAX3::FOXO1 protein expression level.

To investigate potential mechanisms for modulation of chromatin recognition by PAX3::FOXO1 dosage, we identified motifs present in the dosage-specific and common PAX3::FOXO1 sites using HOMER.^44^ We excluded the 0 and 1000 ng/mL conditions from motif calling given their low numbers of unique sites. To visualize trends, we plotted each identified motif by TF class (bZIP, bHLH, etc), given the similarity of the DNA sequences within each class, and by the p-value generated by HOMER, indicating enrichment of that motif within the analyzed set of PAX3::FOXO1 sites (**Figure 3e, Extended Data Figure 2c, see Methods**). Note that the p-values here indicate relative enrichment within each analyzed set of sites and are less informative comparing across sets, meaning that within each dosage level we can use these comparisons to evaluate significance of binding motifs, and then compare the rankings qualitatively for relative enrichments across our dosage ranges. We observe that bZIP (AP-1) and Runt (Runx) motifs are more prominent at 75 and 150 ng/mL than in the common set. In contrast, there is a lower enrichment of bZIP motifs at 500 ng/mL than in the common set. bHLH motifs are less prominent at 75 ng/mL, with increased enrichment in the common set and at higher dosages. The PAX3::FOXO1 composite paired-homeobox motif increases in enrichment as dosage increases and is also enriched in the common set of sites. Homeobox, TEAD, ETS, and zinc finger motifs are present at a low enrichment level across many or all dosages. We also observed that homeobox motifs are just as frequent as bHLH motifs at 75, 250, 500 ng/mL and in the common set of sites (**Extended Data Figure 2d**), despite their lower enrichment in **Figure 3e** due to the high number of homeobox motifs through the genome. We also found highly similar trends at PAX3::FOXO1 sites occupied in two or three adjacent dosages (**Extended Data Figure 2e-h**) supporting the generalizability of our observations.

We hypothesize that the bZIP motifs observed either represent partial PAX3::FOXO1 motif recognition or AP-1 co-binding, present at low dosages. The PAX3::FOXO1 composite and homeobox motifs observed likely reflect PAX3::FOXO1: DNA binding at high dosage. The other motifs observed may represent co-factors of PAX3::FOXO1 or uncharacterized binding preferences of PAX3::FOXO1. Runt (Runx) family co-factors are preferred at lower PAX3::FOXO1 dosages, while usage of bHLH co-factors increase with increasing PAX3::FOXO1 expression. There are several candidate TFs that are expressed in Dbt/MYCN/iP3F and bind the identified motifs (**Supplemental Table 1**). These differences in PAX3::FOXO1 site utilization presumably direct gene expression patterns, leading to different proliferative & oncogenic outcomes driven from distinct epigenetic mechanisms.

### Gene expression is modulated through multiple modes of PAX3::FOXO1-mediated gene regulation and is dosage-dependent

We sought to connect the chromatin-based dose-response effect observed above and the oncogenic and proliferative differences discussed in **Figure 1** by examining gene regulation. We assessed dosage-specific gene expression trends by sequencing RNA from Dbt/MYCN/iP3F compared to an empty vector control (Dbt/MYCN/iEV) exposed to 0-1000 ng/mL doxycycline. Principal component analysis indicated that gene expression is highly similar in Dbt/MYCN/iEV regardless of doxycycline concentration, while in Dbt/MYCN/iP3F samples separate based on doxycycline concentration (**Extended Data Figure 3a**). To identify major trends in gene expression, we performed a gene set variation analysis (GSVA) in Dbt/MYCN/iP3F (**Figure 4a**).^45^ Many cancer-relevant gene sets change significantly with respect to PAX3::FOXO1 dosage. The most prominent gene sets that increase in expression with dosage include S phase-(“E2F targets”) and M phase-specific genes (“G2M checkpoint”, “Mitotic spindle”). Gene sets that decrease in expression with dosage include “Epithelial mesenchymal transition” (EMT) and “Hypoxia.” Interestingly, myogenic genes (“Myogenesis”) have high expression at the intermediate dosage of 150 ng/mL. We do not observe strong doxycycline-dependent trends in Dbt/MYCN/iEV (**Extended Data Figure 3b**). These findings suggest that PAX3::FOXO1 dosage has a strong impact on gene regulation, either through direct DNA binding, indirect mechanisms, or both.

**Figure 4.**
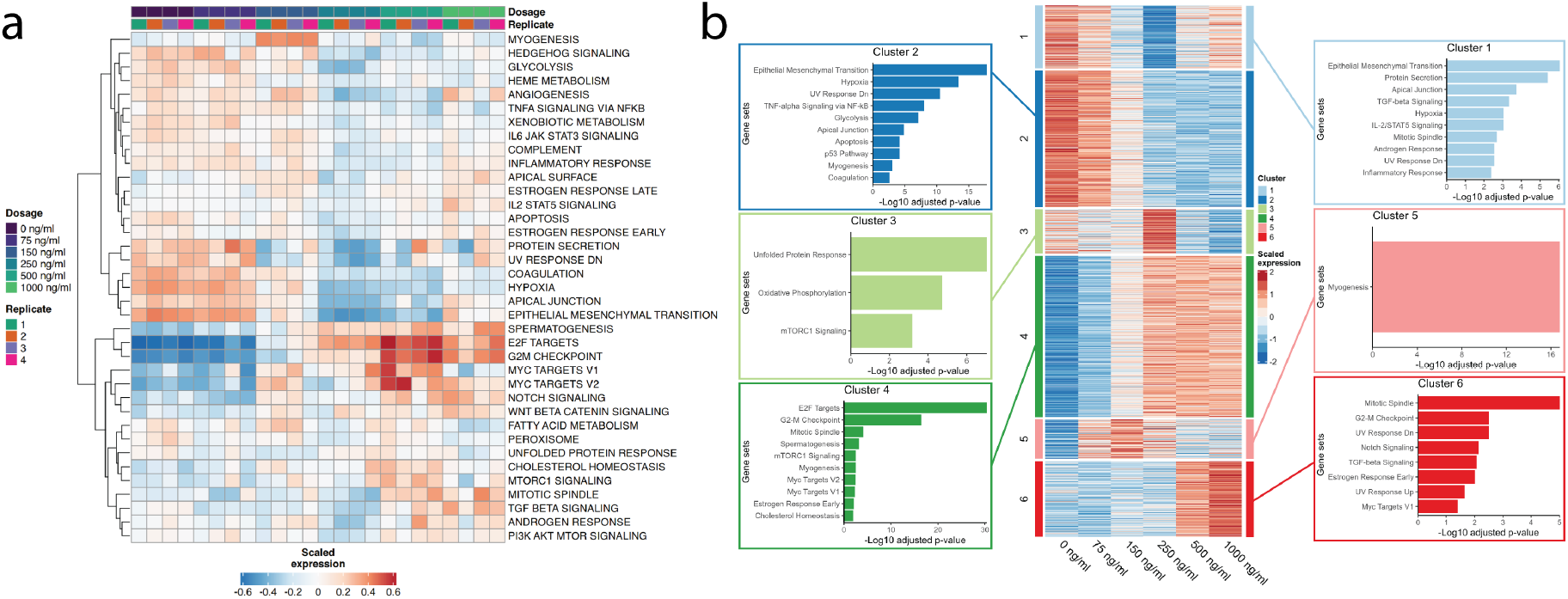
PAX3::FOXO1 dosage regulates expression of oncogenic and developmental pathways. **(a)** Heatmap of scaled expression of gene sets (MSigDB Hallmark) at a range of doxycycline concentrations, from gene set variation analysis of bulk RNA-seq from Dbt/MYCN/iP3F. Replicates are plotted separately and rows are ordered by hierarchical clustering. (**b)** Center: Heatmap of k-means clustered scaled gene expression of differentially enriched genes within 50 Kb of a PAX3::FOXO1 peak. Left, right: Enriched gene sets (from MSigDB Hallmark) in each cluster of genes.

To investigate dosage-dependent gene regulation tied directly to PAX3::FOXO1 DNA binding, we compared the differentially enriched genes identified to the genes within 50 Kb of PAX3::FOXO1 peaks (see **Figure 3**). We applied k-means clustering to expression of the overlapping genes, likely to be directly regulated by PAX3::FOXO1 (**Figure 4b**, center). We obtained the gene sets enriched in each cluster using EnrichR (**Figure 4b** left, right).^46^ Genes decreasing in expression with increasing PAX3::FOXO1 dosage are enriched in EMT, hypoxia, and TNF-alpha signaling gene sets, among other terms. Genes increasing in expression with dosage again include S phase and M phase gene sets. Some M phase gene expression patterns (“mitotic spindle,” “G2M checkpoint”) appear to be highly expressed specifically at the highest doxycycline dosages. We again find that myogenic gene expression is maximized at 150 ng/mL. At 250 ng/mL, genes involved in the unfolded protein response, oxidative phosphorylation, and mTORC signaling have high expression. We also observe a cluster of genes decreasing in expression to 250 ng/mL and then increasing with dosage, including genes involved in EMT, protein secretion, and apical junctions.

The gene expression trends associated with direct PAX3::FOXO1 DNA binding and regulation lead us to hypothesize that PAX3::FOXO1 may act as both a negative and positive regulator of gene expression depending on context. We asked whether trends in DNA motifs might be associated with these different regulatory roles across dosages. Motif enrichment and frequency was highly consistent across all clusters from **Figure 4b**, with the major determinant being PAX3::FOXO1’s expression level, as opposed to its localized gene targets (**Extended Data Figure 3c**,**d**). This indicates that the observed gene expression patterns are not tied to distinct motifs changing in a locus-specific manner. Rather, these data suggest a separation of function between PAX3::FOXO1’s DNA binding capabilities and its recruitment of gene regulatory factors to modify gene expression, which may indeed be locus-specific, based on the adjacent regulatory DNA sequences proximal to a PAX3::FOXO1 binding motif. The mechanisms through which PAX3::FOXO1 directs positive or negative gene regulation will be important to explore in future studies, as well as the combinatorial logic of its molecular recognition with secondary factors at discrete targeting loci. Overall, pathways with gene expression regulated directly by PAX3::FOXO1 include cell cycle pathways and EMT. While myogenic genes may increase in expression at intermediate dosages, many signaling pathways appear to have intermediate expression at intermediate PAX3::FOXO1 dosages.

### PAX3::FOXO1 dosage modulates S phase progression and restoration of chromatin accessibility after replication

Genes tied to the S phase of the cell cycle are highly expressed with high PAX3::FOXO1 expression in our bulk and single cell gene expression datasets, motivating us to investigate S phase progression in the presence of PAX3::FOXO1 at varying dosages. In Dbt/MYCN/iP3F, we quantified the proportion of cells in S phase by measuring incorporation of the thymidine analog EdU (5-Ethynyl-2’-deoxyuridine) (**Figure 5a**). The proportion of S phase cells increases significantly at 500 ng/mL. This aligns with our observation that high PAX3::FOXO1 expression accompanies increased expression of E2F targets and the proportion of single cells expressing S phase genes (**Figure 2c, Figure 4a**). Given the decreased proliferation we observe with increased dosage that precludes faster cycling **(Figure 1e)**, the increased proportion of S phase cells at 500 ng/mL may indicate a prolonged S phase, potentially induced by dosage-mediated transcriptional regulation of replication proteins by PAX3::FOXO1 that could alter various aspects of replication to modify S phase duration.^47,48^ In summary, we hypothesize that the higher percentage of S phase cell specific gene expression signatures indicates failure to progress properly through the cell cycle.

**Figure 5.**
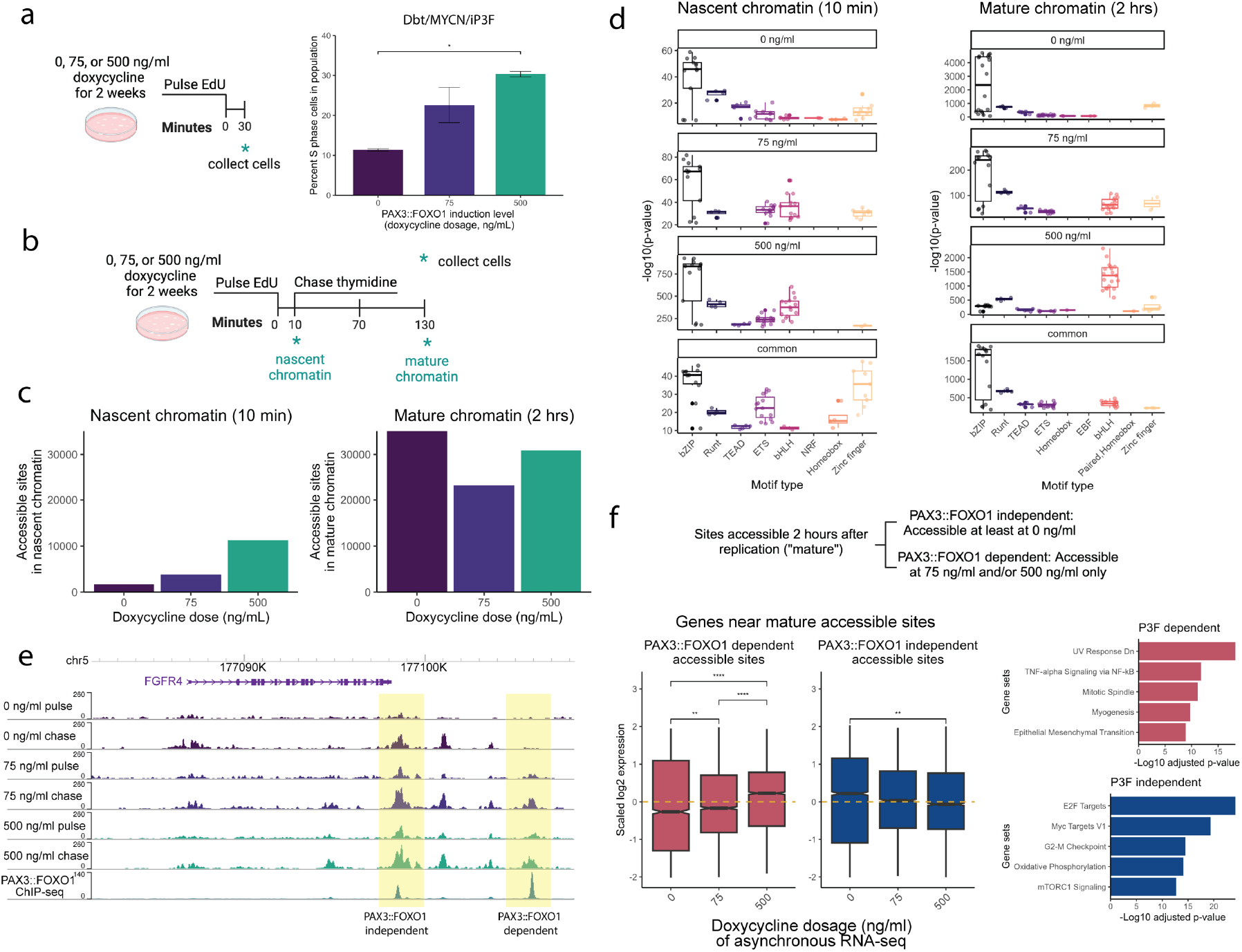
PAX3::FOXO1 dosage perturbs S phase progression and restoration of chromatin accessibility after replication. **(a)** (Left) Schematic for EdU labeling and (right) percent S phase (EdU+) cells in Dbt/MYCN/iP3F by doxycycline concentration at endpoint of a two-week colony formation assay, determined by flow cytometry. **(b)** Schematic for repli-ATAC-seq pulse/chase time course in Dbt/MYCN/iP3F. **(c)** Number of significant repli-ATAC-seq peaks in nascent or mature chromatin by doxycycline concentration. **(d)** Motif enrichment within unique repli-ATAC-seq peaks for each doxycycline concentration and within common peaks across doxycycline concentrations in nascent or mature chromatin. Each motif found is plotted by the - log10 p-value and motif family determined by HOMER. **(e)** Normalized repli-ATAC-seq signal at *FGFR4* in Dbt/MYCN/iP3F. PAX3::FOXO1 ChIP-seq (150 ng/ml) from Figure 3a. **(f)** Top: Scheme for determining PAX3::FOXO1 independent or dependent repli-ATAC-seq sites. Left: Gene expression (asynchronous, from Figure 4) of nearest genes to PAX3::FOXO1-independent or -dependent accessible sites by doxycycline concentration in maturing chromatin. ANOVA with Tukey post-hoc test was used to compare across doxycycline concentrations, ** indicates p ≤ 0.01, *** p ≤ 0.001, **** p ≤ 0.0001. Right: Enriched gene sets (from MSigDB Hallmark) within PAX3::FOXO1-independent or -dependent genes in maturing chromatin.

Given that PAX3::FOXO1 has pioneer function and may transcriptionally modulate S phase progression, we wondered if it could also bind S phase chromatin to modulate the epigenome in a dosage-based manner. We utilized the factor-agnostic approach repli-ATAC-seq to query chromatin behind the replication fork.^49,50^ We used a 10 minute EdU pulse to label replicating DNA in asynchronous Dbt/MYCN/iP3F cells grown in a colony formation assay at 0, 75, or 500 ng/mL of doxycycline for two weeks. We immediately collected cells to obtain nascent chromatin or stopped EdU incorporation with a thymidine chase, collecting mature/maturing chromatin 2 hours later (**Figure 5b**). We isolated EdU-labeled accessible DNA for sequencing to assess nascent and mature chromatin accessibility.

We determined the number of consensus, or reproducible, nascent and mature accessible peaks across replicates in our repliATAC-seq data with respect to PAX3::FOXO1 dosage (**Figure 5c**). In nascent chromatin, we observed a low number of consensus accessible sites, reflecting the variable state of chromatin across cells following replication. We observed 1685 accessible sites without PAX3::FOXO1 induction, increasing to 11,237 sites with high PAX3::FOXO1 dosage. All dosages in mature chromatin have more consensus accessible sites, ranging from 23,290 to 35,066, than in nascent chromatin. In nascent chromatin, the sites unique to each dosage show a similar distribution to the total number of sites at each dosage (**Extended Data Figure 4a**). PAX3::FOXO1 appears to directly or indirectly increase population-level patterns of accessibility within nascent chromatin, which we hypothesize is due to the stabilization of nascent accessibility. We next assessed which types of genomic sites were most accessible (**Extended Data Figure 4b**). In nascent and mature chromatin most sites are in promoters, introns, and intergenic regions regardless of dosage.

To identify TFs participating in chromatin maturation across PAX3::FOXO1 dosages, we analyzed the motifs present at sites unique to each dosage and common across nascent or mature accessible sites (**Figure 5d, Extended Data Figure 4c, see Methods**). While significance is plotted to enable quantitative comparisons within datasets for motif classes at distinct dosages, qualitative comparisons are also possible to evaluate motif enrichment differences across dosages. In nascent chromatin, bZIP (AP-1) motifs are highly enriched across all dosages. At 75 and 500 ng/mL, bHLH motifs are more enriched than at 0 ng/mL. In mature chromatin, bZIP motifs are highly enriched at 0 and 75 ng/mL, but not at 500 ng/mL. bHLH motifs again are enriched at 75 and 500 ng/mL. The increase in enrichment of bHLH motifs in nascent and mature accessible sites with PAX3::FOXO1 expression is reminiscent of the bHLH motifs at PAX3::FOXO1 sites observed in asynchronous cells (**Figure 3e**), and given this we hypothesize that the increase in bHLH motifs at nascent and mature accessible sites may reflect PAX3::FOXO1 chromatin binding.

PAX3::FOXO1’s influence on chromatin maturation is further demonstrated at the *FGFR4* locus. One of the two PAX3::FOXO1 binding sites downstream of *FGFR4* gains accessibility regardless of PAX3::FOXO1 dosage (“PAX3::FOXO1-independent”), while the other site is only accessible in nascent and mature S phase chromatin at 75 and 500 ng/mL (“PAX3::FOXO1-dependent”, **Figure 5e**). To determine if PAX3::FOXO1-dependent accessible sites in nascent and mature chromatin have any association with local gene expression, we categorized nascent and mature accessible sites into groups based on their accessibility in the 0, 75 and/or 500 ng/mL doxycycline conditions (**Figure 5f, Extended Data Figure 4d**). Genes near S phase accessible sites dependent on PAX3::FOXO1 in both nascent and mature chromatin include members of the FP-RMS CRC (MYOD1, PITX3, SOX8, RARA, FOSL2, PKNOX2; **Supplemental Table 2**). Genes near both PAX3::FOXO1-dependent and -independent sites in nascent chromatin are enriched in cell cycle and cell growth gene sets. In contrast, genes near mature PAX3::FOXO1-dependent sites are associated with processes such as TNF alpha signaling,^51^ M phase,^34,38^ myogenesis,^36^ and EMT,^28,52^ while genes near mature independent sites are involved in cell growth and cycling functions. We then examined gene expression at these genes proximal to PAX3::FOXO1-dependent and -independent accessible sites using our interphase gene expression data at 0, 75, and 500 ng/mL from **Figure 4**. In nascent chromatin, we found that genes near accessible sites had increasing interphase RNA expression as dosages increased from 0 and 75 ng/mL to 500 ng/mL, with slightly stronger effects near PAX3::FOXO1-dependent genes (**Extended Data Figure 4d**). In mature chromatin, PAX3::FOXO1-dependent accessible sites are associated with increases in interphase gene expression with dosage, while PAX3::FOXO1-independent accessible sites in mature chromatin are associated with decreases in interphase gene expression with dosage (**Figure 5f**). These observations hint at mechanistic connections between PAX3::FOXO1-dependent accessible sites during chromatin maturation and gene expression patterns that drive FP-RMS.

### PAX3::FOXO1 binding sites gain nascent and mature chromatin accessibility with high PAX3::FOXO1 expression

As the PAX3::FOXO1-dependent accessible sites we observed might be bound directly by PAX3::FOXO1 or be indirectly regulated by PAX3::FOXO1 through downstream factors, we investigated whether PAX3::FOXO1 binding sites have altered nascent accessibility. We utilized PAX3::FOXO1 sites identified in **Figure 3**. We plotted nascent accessibility across PAX3::FOXO1 expression levels at these PAX3::FOXO1 sites (**Figure 6a**, left), and used k-means clustering to identify three clusters with distinct differences in accessible signal. Notably, cluster 3 is composed of sites with low chromatin accessibility at 0 and 75 ng/mL doxycycline. In all clusters, accessibility increases with PAX3::FOXO1 expression, and we hypothesize that this relates to pioneering activity that may be occurring during chromatin maturation. Similar trends are present in mature chromatin (**Figure 6a**, right).

**Figure 6.**
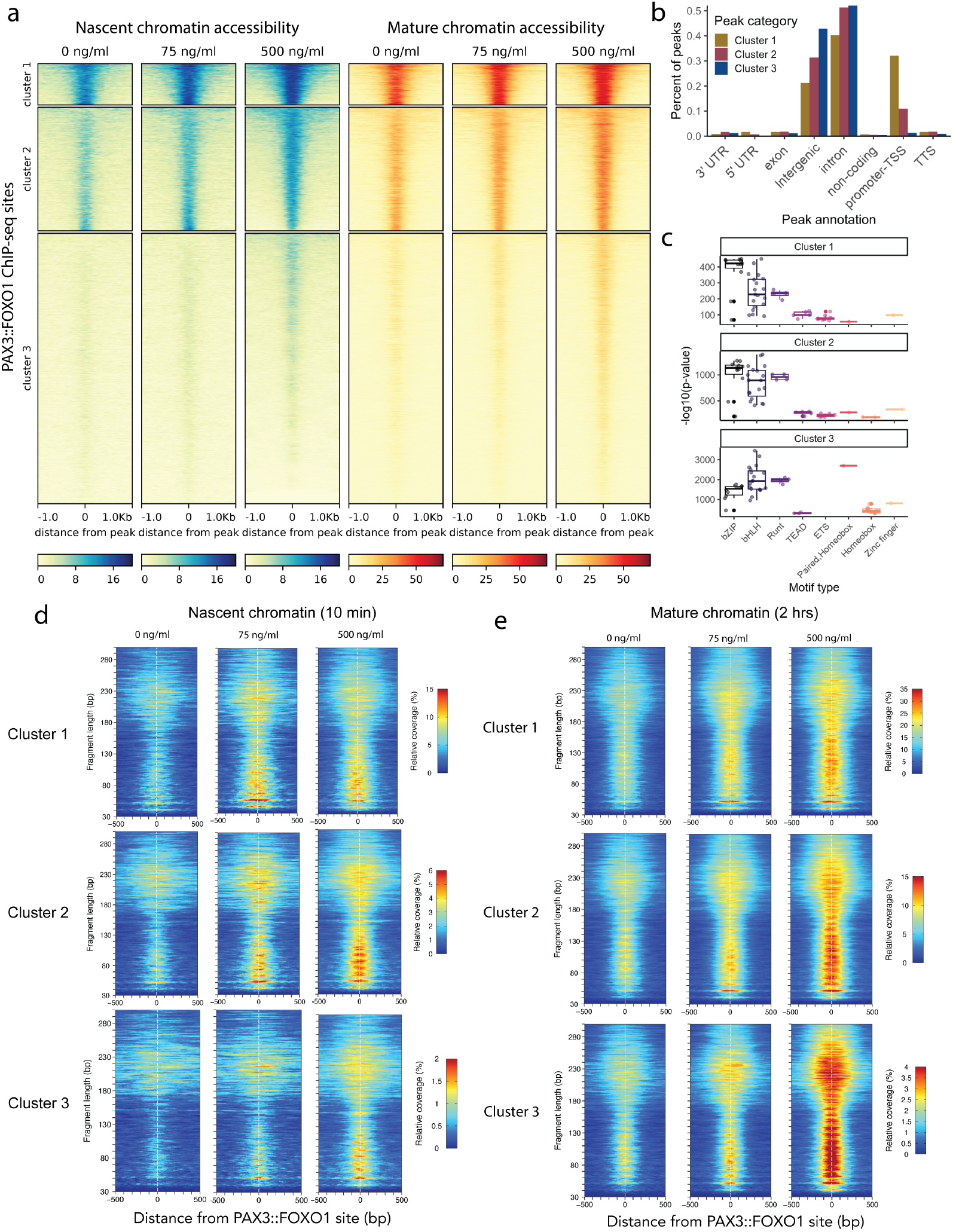
Nascent and mature chromatin accessibility stratifies PAX3::FOXO1 sites. **(a)** k-means clustered repli-ATAC-seq accessible signal at PAX3::FOXO1 ChIP-seq sites (all sites across dosages, from Figure 3) in nascent and mature chromatin by doxycycline dosage. **(b)** Distribution of PAX3::FOXO1 sites in each genome annotation category, by cluster. **(c)** Motif enrichment at PAX3::FOXO1 sites in each cluster. Each motif found is plotted by the -log10 p-value and motif family determined by HOMER. Nascent chromatin **(d)** and mature chromatin **(e)** repli-ATAC-seq fragment length distribution around PAX3::FOXO1 sites in each cluster across doxycycline concentrations.

To distinguish these clusters of PAX3::FOXO1 sites, we examined the characteristics of each set of sites. All three clusters contain sites located in intergenic and intronic regions. Cluster 1, which is highly accessible, also has approximately 32% of sites located in transcription start sites (**Figure 6b**). We next identified the pathways enriched amongst the nearest genes to the PAX3::FOXO1 sites in each cluster (**Extended Data Figure 5a**). Genes involved in hypoxia, myogenesis, TNF alpha signaling, mitotic spindle function, and EMT were enriched in all clusters, indicating that many processes may be impacted by PAX3::FOXO1’s influence in chromatin replication. Cluster 3 contains a higher proportion of 500 ng/ml dosage-specific PAX3::FOXO1 sites (23%) (**Extended Data Figure 5b**), aligning with the accessibility observed in cluster 3 at 500 ng/mL. Finally, we performed a motif analysis on each cluster of PAX3::FOXO1 sites (**Figure 6c, Extended Data Figure 5c**). bHLH motifs are enriched across all clusters as in **Figure 5d**. In cluster 3, bZIP motifs are less enriched while the PAX3::FOXO1 composite motif is more enriched in comparison to clusters 1 and 2. In summary, cluster 1 sites are highly accessible regardless of PAX3::FOXO1 expression, reflecting their frequent proximity to transcribed regions of the genome. Cluster 2 sites are likely farther from transcribed genes but have accessibility regardless of PAX3::FOXO1 expression, while cluster 3 sites are less accessible throughout chromatin maturation except in the presence of high PAX3::FOXO1 expression. Cluster 3 sites are also often occupied at high PAX3::FOXO1 dosage. These observations provide rationale for further studies to investigate mechanisms of pioneering during chromatin replication and maturation, and suggest that PAX3::FOXO1 modulates S phase chromatin accessibility at known PAX3::FOXO1 sites in a dosage-dependent manner.

To investigate whether the increase in accessibility we observed across these clusters represented nucleosome phasing and/or transcription factor occupancy, we visualized the fragment size distribution in each cluster in nascent and mature chromatin (**Figure 6d**,**e, Extended Data Figure 5d**,**e**).^53^ In nascent chromatin, accessible signal across all fragment sizes increases similarly with both low and high PAX3::FOXO1 expression in clusters 1 and 2. Cluster 3 has a greater increase in accessible fragments at 500 ng/mL compared to 75 ng/ml. In mature chromatin, accessible signal increases most with high PAX3::FOXO1 dosage in all three clusters, though this is again greater in cluster 3. PAX3::FOXO1 (or a PAX3::FOXO1-regulated pioneer) may be involved in both nucleosome phasing and eviction at these sites in a dosage-dependent manner. We propose that PAX3::FOXO1 plays a role in modulating chromatin maturation immediately after fork passage, establishing epigenetic memory in FP-RMS and altering epigenetic memory in the originating cell.

## Discussion

We have demonstrated that PAX3::FOXO1 expression level, or dosage, modulates oncogenic character and also induces dosage-specific chromatin recognition. In our model, an intermediate dosage of PAX3::FOXO1 optimizes focus formation and thus oncogenic potential. However, PAX3::FOXO1 dosage likely varies across a population of tumor cells, and other PAX3::FOXO1 dosages lend insight into how PAX3::FOXO1 may facilitate a variety of cellular phenotypes via dosage-dependent gene regulation. ***Low PAX3::FOXO1 expression*** is associated with low levels of focus formation and unhindered proliferation. bZIP motifs have sequence similarity to part of the PAX3::FOXO1 composite motif, so these motifs could represent an AP-1 family co-factor or partial motif recognition by PAX3::FOXO1 as part of its pioneer function at low dosages. High expression of gene regulatory programs including EMT, hypoxia, and angiogenesis is retained. Alongside this, we observe dosage-specific PAX3::FOXO1 site usage based on the motifs present. The combinatorial logic of gene regulation at PAX3::FOXO1 sites may vary with dosage, requiring different cooperating factors, motifs, and adjacent PAX3::FOXO1 sites.^54,55^ It will be intriguing to resolve these relationships in future studies.

***Intermediate PAX3::FOXO1 expression*** maximizes focus formation and cells exhibit proliferation levels in between those found at low and high dosages. The number of PAX3::FOXO1 binding sites is also maximized, and bHLH motifs may represent myogenic co-factors (MYOG, MYOD1) relevant at intermediate dosages and greater. A subset of gene expression programs highly expressed at low and high dosages are moderately expressed with intermediate PAX3::FOXO1 expression, while myogenic and metabolic gene expression increases.

Finally, ***high PAX3::FOXO1 expression*** leads to decreases in proliferation and focus formation. Higher expression of cell cycle gene expression suggests that either (i) uncontrolled cycling might result if cells escaped the suppression of high PAX3::FOXO1 dosage, or (ii) increased cell cycle gene expression may lead to induced growth suppression, or a combination of these mechanisms.^56,57^ PAX3::FOXO1 exhibits increased binding to its complex composite paired-homeobox motif at high dosage, which may indicate another mode of pioneer motif recognition. There are clear dosage-dependent trends in both PAX3::FOXO1 motif recognition and potential co-factor motifs across dosages suggesting that PAX3::FOXO1 dosage may modulate the mode of its pioneering activity. We speculate that low and high PAX3::FOXO1 expression levels could have roles to play throughout tumor initiation and progression; for example, low PAX3::FOXO1 expression has been shown to favor migration.^58^ Prior studies demonstrate that PAX3::FOXO1 gene and protein expression varies across tumor models and across individual cells.^30,59^ We hypothesize that variegated PAX3::FOXO1 expression, both at the cell population level between RMS tumors, and in the context of cell-to-cell variability in expression within a tumor, provides selective advantages during sarcomagenesis given the spectrum of potentially advantageous characteristics observed across dosages.

Our findings may be relevant for other oncogenic fusion TFs. Though the dosage-specific gene-regulatory roles of other oncogenic TFs may not be precisely the same, similar principles may hold. Aspects of our data fit a Goldilocks principle, defined as observations that increase with PAX3::FOXO1 dosage until an intermediate inflection point and then decrease with increasing dosage. The number of PAX3::FOXO1 binding sites, the expression of some myogenic genes, and focus formation with dosage fit this model. In contrast, other gene expression patterns we observed, proliferation in bulk culture, and PAX3::FOXO1 binding sites in nascent chromatin instead increase or decrease across all dosages tested in this study. Related observations have been made in Ewing sarcoma. Low expression of EWSR1::FLI1 favors migration,^60^ analogous to prior observations of PAX3::FOXO1 and the higher expression of EMT-related genes observed at low PAX3::FOXO1 dosages in this study.^28^ However, high expression of EWSR1::FLI1 can support high proliferation, unlike the observations in our current study.^60^ High expression of EWSR1::FLI1 obtainable in the absence of TRIM8 is toxic to cells,^61^ analogous to the growth suppression observed in this study at high PAX3::FOXO1 dosage. Even though these factors and their functions are not identical, dosage of the fusion TF matters for oncogenic character in both FP-RMS and Ewing sarcoma. Future work on PAX3::FOXO1 should explore whether a targetable regulator of PAX3::FOXO1 dosage can be identified analogous to TRIM8. Other mutationally quiet cancers with TF drivers may also benefit from mechanistic studies of dosage.

PAX3::FOXO1 appears to facilitate heterogeneity of gene expression in human myoblasts following prolonged expression within a system allowing for environmental heterogeneity. Do TFs in general have the ability to facilitate heterogeneity? This would not necessarily be favorable, given that maintaining a stable cell identity is necessary for tissue specification and function. However, multiple cell identities have been identified within FP-RMS patient-derived xenografts, and PAX3::FOXO1 may promote the establishment of these populations.^33,34,37–39^ The ability to facilitate heterogeneity may be a feature of oncogenic TFs, and understanding this will be important context for any new therapies developed to target these factors. Some of the populations we observed in Dbt/MYCN/iP3F are analogous to identities found in RMS tumors, supporting the utility of our model system. The expression of CRTFs also varies across single cells, as opposed to one unifying CRC across all cells. The ability of CRTFs to act concordantly at the same regulatory elements is dependent upon their co-expression within a cell, and there is not one particular cell state accompanying each PAX3::FOXO1 expression level we tested. The FP-RMS CRC may be an emergent phenomenon more readily observed in bulk cells. Alternatively, we may have captured an intermediate state on the trajectory from myoblast to FP-RMS cell that has yet to establish the full CRC. The extent to which this is generalizable to other models and cancers remains an open question, and the role of the CRC in single cells in FP-RMS and other cancers should be a subject of future studies.

Our studies also implicate multiple gene-regulatory modes for PAX3::FOXO1. Genes near PAX3::FOXO1 sites can either increase or decrease in expression with PAX3::FOXO1 dosage. In addition to PAX3::FOXO1’s long-established role as a transcriptional activator,^62,63^ it may act as a repressor at certain sites. Similar bimodal function has been observed for other TFs.^64^ A recent study of TF effector domains identified repressive and activating domains in regions of FOXO1 preserved in PAX3::FOXO1.^65^ We predict that the intrinsically disordered C terminal region of FOXO1 may interact with repressive or activating factors depending on local chromatin context. If PAX3::FOXO1 has a role in altering gene-regulatory looping, that might also contribute to bimodal regulation. Future investigations may provide insight into a spectrum of PAX3::FOXO1 functions.

We hypothesize that the loss of epigenetic memory of the initial cell state in FP-RMS is mechanistically linked to PAX3::FOXO1’s function to promote transcriptional heterogeneity. Chromatin re-establishment after replication in a cancer context is severely understudied, and we have sought to apply knowledge of chromatin replication and maturation to the context of pediatric sarcoma. Prior studies have examined subnucleosomal footprints likely established by TFs and confirmed that TFs can bind rapidly behind the fork.^49,66,67^ We have demonstrated that a fusion TF influences accessibility rapidly behind the replication fork in a dosage-dependent manner. PAX3::FOXO1 may directly bind to nascent and maturing chromatin. We also speculate that PAX3::FOXO1’s pioneering function may have a role in nascent chromatin recognition, given the lack of consistent accessibility behind the replication fork.^16^ Alternatively, other factors regulated by PAX3::FOXO1 could establish the observed PAX3::FOXO1-dependent accessible sites. PAX3::FOXO1-dependent sites present 2 hours after replication during maturation also appear to influence transcription, suggesting a mechanism for PAX3::FOXO1 to alter gene regulation through binding to newly replicated chromatin. Future studies of PAX3::FOXO1, its co-factors, and histone modifications in replicated chromatin will help refine this model of chromatin replication in FP-RMS, which may serve a fundamental mechanism for propagating epigenetic information.

## Study limitations

We note that PAX3::FOXO1 protein levels may vary across cells in a population exposed to the same concentration of doxycycline. Given the PAX3::FOXO1 dosage sensitivity of our system, we posit that despite this potential variability, PAX3::FOXO1 expression on average is distinct between dosages. This work proposes mechanisms for PAX3::FOXO1’s chromatin recognition which remain to be confirmed through future biochemical work. The gene-regulatory outcomes of PAX3::FOXO1 dosage are diverse and thus could not all be mechanistically investigated here. Our use of repli-ATAC-seq yielded factor-agnostic information which does not confirm which factors bind to each accessible site. The focus of this work is on dosage of PAX3::FOXO1 in a context resembling tumorigenesis and does not address other roles for PAX3::FOXO1. Our Dbt/MYCN/iP3F system offers one model for PAX3::FOXO1 dosage, and additional aspects of PAX3::FOXO1 dosage may remain to be uncovered in other cellular contexts.

## Acknowledgements

The authors thank current and former members of the Stanton and Barr groups for support and feedback. We are grateful to Ryan Roberts, Rachid Drissi, and the OSU/NCH epigenetics community for suggestions and advice. RAH thanks Timothy Cripe, Genevieve Kendall, and Christine Cucinotta for key comments and suggestions. We thank Li-Chun Tu and Sonja Chen for comments on the manuscript. RAH is supported by NCI T32CA269052 and a CancerFreeKids New Idea Award. BZS is grateful to American Cancer Society (RSG-23-1021178-01-DMC), St. Baldrick’s Foundation (Career Development Award), National Institutes of Health (R01GM144601, 1R01HL166520 - 01A1), and intramural funding from Nationwide Children’s Hospital for supporting this work. FGB is supported by the intramural program of the National Cancer Institute and the Joanna McAfee Childhood Cancer Foundation. Figures 1a and 5a, b, and f were created in BioRender.

## Methods

### Cell culture

All cell lines were cultured in F-10 medium for population growth, colony growth and focus formation as previously described.^29,30^ For doxycycline induction, cells were plated in fresh medium and allowed to grow for 24 hours, and then the medium was replaced with fresh medium containing the desired concentration of doxycycline (Sigma D9891) and then changed at 2-3 day intervals.

### Western blotting

Cells were resuspended in PBS containing protease inhibitors (Active Motif #37491) and then an equivalent volume of SDS loading buffer (Quality Biological 351082661). They were then incubated at 95°C for 10 minutes to obtain whole cell lysates. Samples were run on a NuPAGE 4-12% BisTris gel and transferred to a nitrocellulose membrane either at 4°C/30V overnight or for 1 hour/100V at room temperature. Membranes were incubated with primary antibodies (1:500 FOXO1 (Cell Signaling, 2880S), 1:1000 TBP (Cell Signaling, 44059S)) in 5%/TBST (0.1% v/v Tween-20) either for 2 hours at room temperature or overnight at 4°C. Membranes were then incubated for 1 hour with HRP-conjugated secondary antibody (Invitrogen #31460), incubated with SuperSignal West Pico PLUS Chemiluminescnent substrate (ThermoFisher #34580) and imaged with a LiCor C-DiGit Blot Scanner and Image Studio software (v5.5).

### Proliferation assays

Experiments were performed in a Incucyte S3 Live Cell Analysis System. Cells were seeded into 96-well plates and allowed to grow in a cell culture incubator for 24 hours. Media containing doxycycline at 2X the final concentration was added to each well. Plates were scanned in the Incucyte S3 using the Phase channel for 60 hours. Image analysis to quantify confluence was performed in Incucyte Live Cell Analysis 2023A software and data was visualized using the ggplot package in R.^68^

### Focus formation and colony formation assays

Focus formation experiments and colony formation assays were performed as previously described.^59,69^

### Single cell RNA-sequencing & analysis

Cells were collected, viably frozen, and sent out for 10X Genomics Chromium 3’ single-cell RNA-sequencing by Azenta Life Sciences. Data was aligned using the Cell Ranger pipeline. Data was analyzed using the Seurat package (v5.0.1) in R.^70^ After quality control filtering, the data was log normalized using NormalizeData and cell cycle annotations were assigned using CellCycleScoring and the file cycle.rda available from Seurat.^71^ Data was scaled using ScaleData and batch-corrected using Harmony.^72^ Data was clustered at a resolution determined using clustree.^73^ UMAP projections were obtained with RunUMAP. Progenitor, proliferative, and differentiated scores were assigned using AddModuleScore and the gene sets from Danielli et al 2024.^34^ Significant markers for each cluster were obtained using FindAllMarkers with logfc.threshold= 0.25, min.pct = 0.1, and min.diff.pct=0.1. Markers were analyzed using EnrichR and fgsea to determine enriched gene expression programs.^46,74^ To obtain clusters with significantly different expression patterns, the initial set of clusters were subclustered or merged as needed manually. Progenitor, proliferative, and differentiated scores for each cluster in addition of gene set enrichment results were utilized for cluster annotation. These analyses were performed separately for the short and long timepoint samples. Visualizations were performed using the ggplot R package in addition to Seurat.

### ChIP-sequencing & analysis

Chromatin immunoprecipitation and sequencing was performed as previously described with 2 biological replicates.^24^ Reads were aligned to hg38 using the ENCODE ChIP-seq pipeline (v2.1.2). Peaks were called within the pipeline by macs2. Unique and overlapping peak categories were determined with a custom R script and the GenomicRanges package.^75^ The WashU Epigenome Browser was used for data visualization.^76^ HOMER (v4.11) was used to called motifs and annotate ChIP-seq peaks.^44^ Motif enrichment was plotted in R by first filtering TF motifs identified by the expressed TFs in the Dbt system. Motifs were then plotted by the - log10 p-value and motif category/family obtained from HOMER. Because the log 10 p-value from HOMER analyses is influenced by the number of sites upon which motifs are called,^44^ we chose to make the y-axes of these plots scaled based on the maximum log10 p-value within each set of peaks so that relative comparisons of the most significant motifs in each dataset are emphasized.

### RNA-sequencing & analysis

Paired-end RNA sequencing was performed on the NovaSeq platform (Illumina) at Novogene, and sequencing data were processed for alignment using HISAT2 and TOPHAT2. Principal component analysis and differentially expressed gene identification were performed with DESeq2 in R.^77^ Scaled log2 gene expression was used as input for gene set variation analysis (GSVA) and for heatmaps of gene expression. GSVA was performed using the GSVA and limma packages in R.^45,78^ Heatmaps were generated using the pheatmap and ComplexHeatmap packages in R.^79,80^ enrichR was used for gene set analyses.^46^

### Flow cytometry

EdU/DAPI labeling and flow cytometry were performed as previously described with modifications.^81^ Cells were seeded and once they reached 50-80% confluence, EdU was added at a final concentration of 20 nM. Cells were incubated for 30 minutes and then collected by trypsinization. Cells were washed in PBS and then resuspended in fresh 70% ethanol with vortexing. Cells were fixed at -20°C for overnight or longer. Cells were then resuspended and washed in 1% FBS/PBS. To visualize EdU, a Click reaction was performed by resuspending cells in 2 mM CuSO4, 20 mg/ml ascorbate, and 10 uM sulfo-cyanine azide and incubating for 30 minutes at room temperature. Cells were then washed in FBS/PBS, resuspended in 1:1000 DAPI (1 mg/ml stock, ThermoScientific #62248) FBS/PBS, and analyzed on a BD Fortessa. Data analysis was performed in FlowJo (v5.0) and visualized in R with ggplot2.^68^

### repli-ATAC-sequencing & analysis

repli-ATAC-seq was performed in biological replicates according to the published protocol with minor modifications.^50^ Cells were pulsed with 20 nM EdU (final concentration) for 10 minutes. Cells were then collected by trypsinization or washed with PBS and incubated with media containing 10 uM thymidine for a 2 hour chase. A “no EdU” plate of cells was also collected and processed through library preparation as a control. During library preparation, the original ATAC-seq index primers were used with 2X Phusion Master Mix (NEB #M0531L) for the on-bead PCR. The number of PCR cycles to use for each sample was determined using qPCR as in the original ATAC-seq protocol.^82^

Reads were aligned to hg38 using the ENCODE ATAC-seq pipeline (v2.2.2) and peaks were called using macs2 within the pipeline. Unique and overlapping peak categories were determined with a custom R scripts as in ChIP-seq analyses. Motifs were called and peak annotations were determined using the HOMER software package.^44^ Motif enrichment results were plotted with the same approach used for the ChIP-seq data. ATAC-seq bigwigs were obtained with deeptools bamCoverage –normalizeUsing RPGC (v3.3.1).^83^ Heatmaps were generated using deeptools and fragment size distributions were plotted with plot2DO (v1.0).^53^ enrichR was used for gene set analyses.^46^ The WashU Epigenome Browser was used for data visualization.^76^ To analyze the effect of PAX3::FOXO1-dependent and -independent accessible sites on gene expression, the nearest gene annotations of the accessible sites were filtered by expressed genes in the bulk RNA-seq data. Scaled log2 gene expression was then plotted based on the P3F dependence of the corresponding accessible site.

## Data availability

Raw sequencing files from this study will be made available in the Gene Expression Omnibus (GEO) upon publication. Scripts used in data analysis are available at https://github.com/rah60/P3F-dosage-2025.

## Notes

### Competing Interest Statement

The authors have declared no competing interest.

