## Supplemental Figures and Table S1 for "Fusion transcription factor dosage controls cell state in rhabdomyosarcoma"

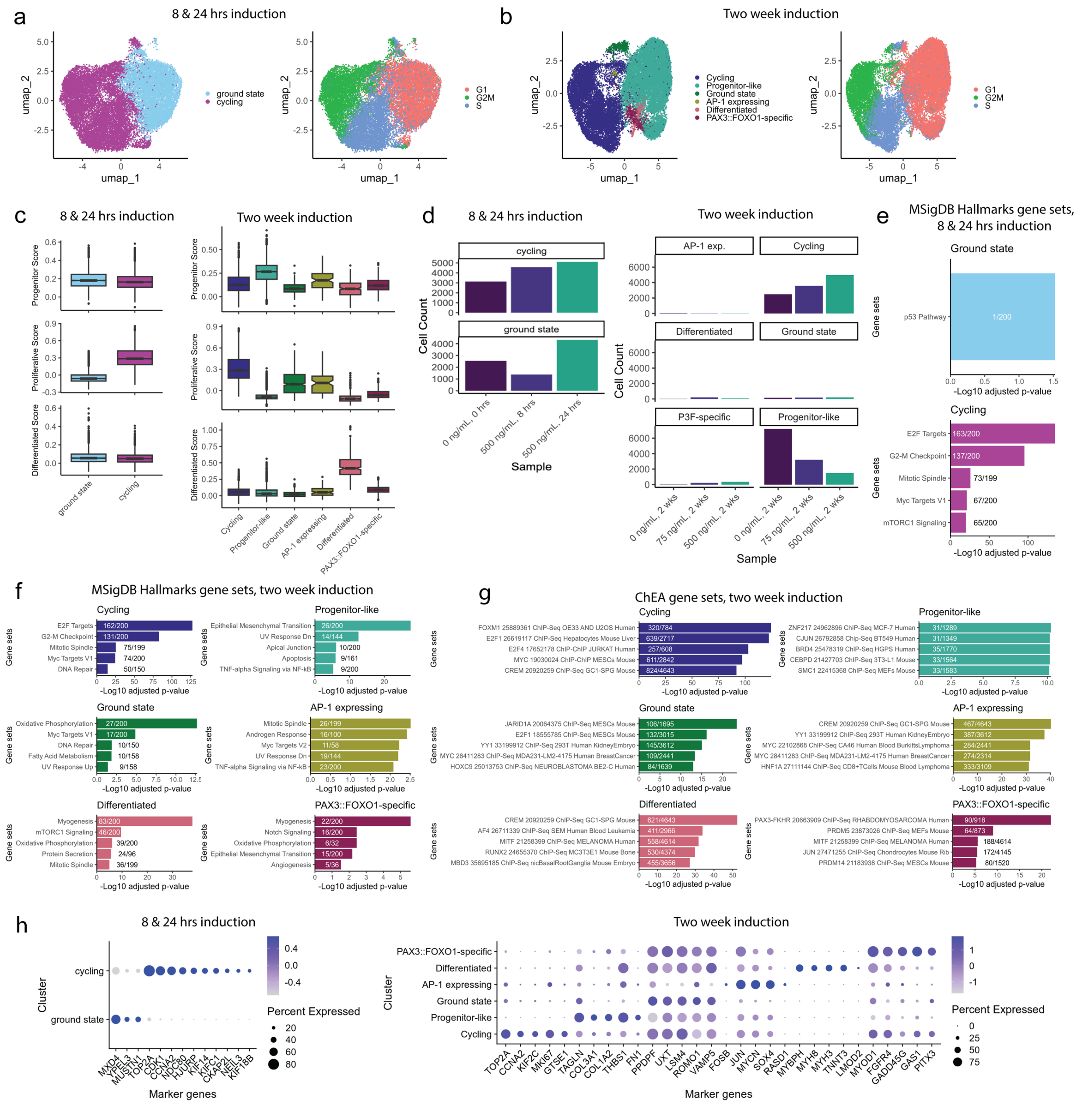

**Extended Data Figure 1. (a)** UMAP projection of Dbt/MYCN/iP3F single cell gene expression after 0, 8, or 24 hours of 500 ng/ml doxycycline induction with clusters labeled (left) and cell cycle state annotations labeled (right). **(b)** UMAP projection of Dbt/MYCN/iP3F single cell gene expression after 2 weeks of doxycycline induction at 0, 75, or 500 ng/ml, with clusters labeled (left) and cell cycle state annotations labeled (right). **(c)** Proliferative, progenitor, and differentiated cell identity scores for Dbt/MYCN/iP3F cells after short or long doxycycline induction, grouped by cluster. Cell identity scores were calculated based on Danielli et al 2024. All boxplots indicate median, first quartile, and third quartile and whiskers indicate 1.5x the inter-quartile range. 8 & 24 hours: ground state n= 8244; cycling n=12827. Two week: Cycling n=11049; progenitor-like n=11925; ground state n= 563; AP-1 expressing n=98; differentiated n=294; PAX3::FOXO1 specific n=572. **(d)** Number of Dbt/MYCN/iP3F cells in each 8 & 24 hr or

2 week induction cluster, grouped by condition. Two week clusters also plotted in Figure 2d. **(e)** Enriched gene sets from MSigDB Hallmark in the differentially expressed genes (“markers”) for each single cell cluster after 0, 8, or 24 hours of 500 ng/ml doxycycline induction. Fraction of genes from each gene set found in single cell markers are indicated on the corresponding bar. **(f)** Enriched gene sets from MSigDB Hallmark in the differentially expressed genes for each single cell cluster after 2 weeks of doxycycline induction at 0, 75, or 500 ng/ml. Fraction of genes from each gene set found in single cell markers are indicated. **(g)** Enriched gene sets from ChEA (ChIP Enrichment Analysis, transcription targets identified by ChIP-based methods) in the differentially expressed genes for each single cell cluster after 2 weeks of doxycycline induction at 0, 75, or 500 ng/ml. Fraction of genes from each gene set found in single cell markers are indicated. **(h)** Marker genes in single cell clusters plotted by scaled expression and percent cells expressing in each cluster in short (left) or long doxycycline induction (right).

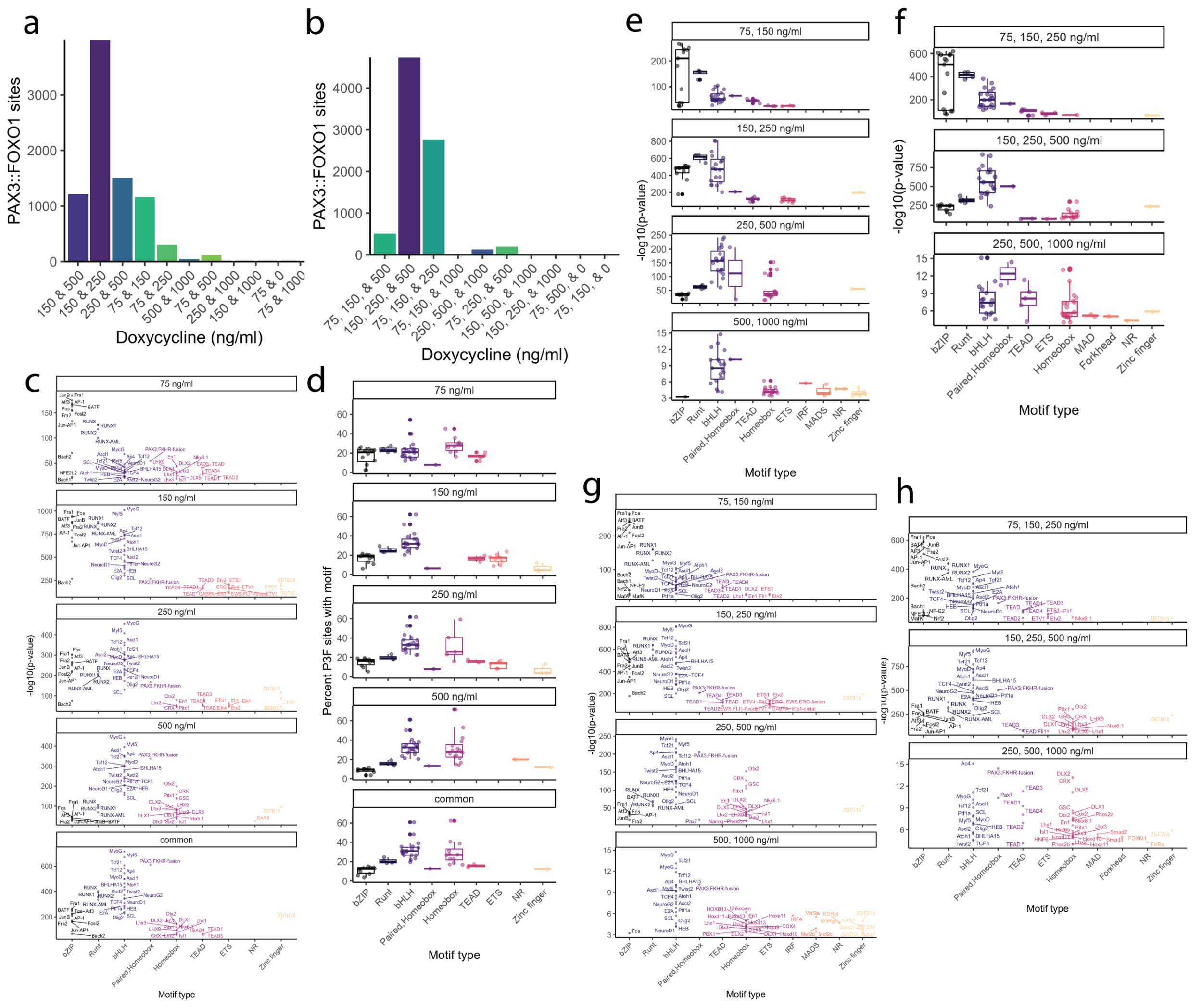


**Extended Data Figure 2. (a)** Number of PAX3::FOXO1 sites occupied across two doxycycline dosages. **(b)** Number of PAX3::FOXO1 sites occupied across three doxycycline dosages. **(c)** Motif enrichment within unique PAX3::FOXO1 peaks to each doxycycline concentration and within common peaks across all doxycycline concentrations. Same as Figure 3e with motif names labeled. Each motif found is plotted by the -log10 p-value and motif family determined by HOMER. All boxplots indicate median, first quartile, and third quartile and whiskers indicate 1.5x the inter-quartile range. **(d)** Percent motif occurrence within unique PAX3::FOXO1 peaks to each doxycycline concentration and within common peaks across all doxycycline concentrations. Each motif found is plotted by the percent of sites containing motif and motif family determined by HOMER. **(e)** Motif enrichment within PAX3::FOXO1 sites occupied across two doxycycline dosages (adjacent dosages only). **(f)** Motif enrichment within PAX3::FOXO1 sites occupied across three doxycycline dosages (adjacent dosages only). **(g)** Same as Extended Data Figure 3e with motif names labeled. **(h)** Same as Extended Data Figure 3f with motif names labeled.

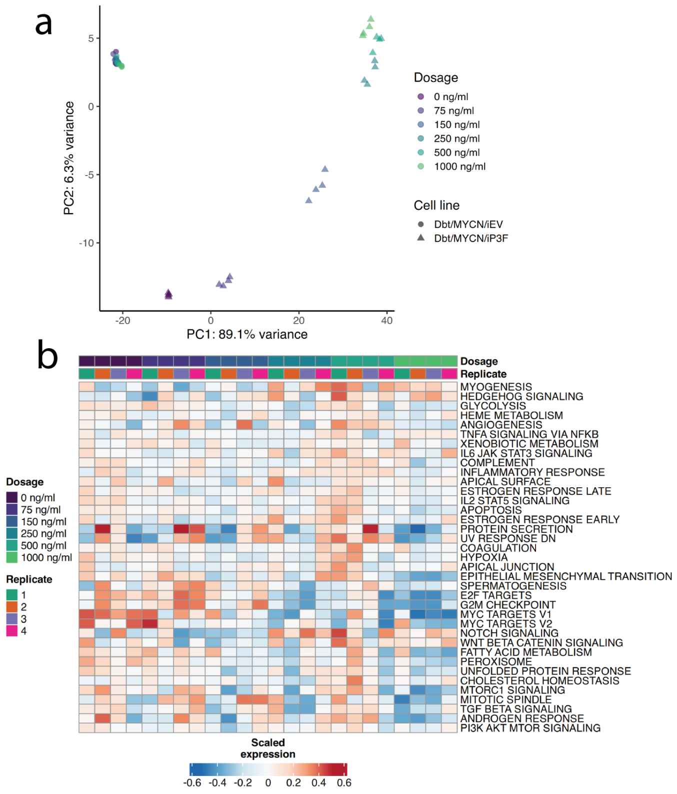


**Extended Data Figure 3.** **(a)** Principal component analysis of bulk RNA-seq samples from Dbt/MYCN/iP3F and Dbt/MYCN/iEV across doxycycline dosages. **(b)** Heatmap of scaled expression of gene sets (MSigDB Hallmark) at a range of doxycycline concentrations, from gene set variation analysis of bulk RNA-seq from Dbt/MYCN/iEV. Replicates plotted are separately and rows are ordered as in Figure 4a. **(c)** Motif enrichment at PAX3::FOXO1 sites by nearest gene expression cluster from Figure 4b. Each motif found is plotted by the -log10 p-value and motif family determined by HOMER. All boxplots indicate median, first quartile, and third quartile and whiskers indicate 1.5x the inter-quartile range. **(d)** Percent motif occurrence at PAX3::FOXO1 sites by nearest gene expression cluster from Figure 4b. Each motif found is plotted by the percent of sites containing motif and motif family determined by HOMER.

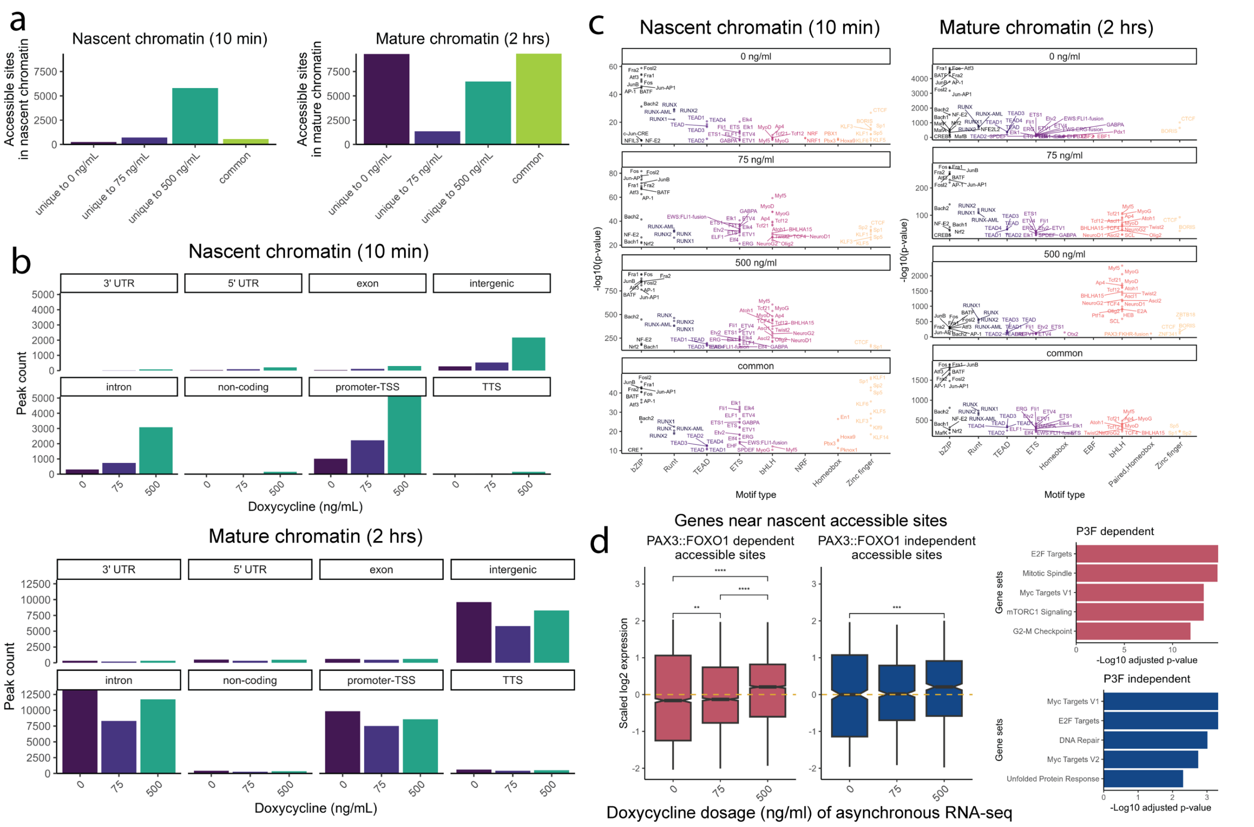


**Extended Data Figure 4. (a)** Number of nascent or mature accessible sites unique to each doxycycline dosage or common across all dosages. **(b)** Distribution of genomic annotations of nascent or mature accessible peaks, by doxycycline dosage. **(c)** Motif enrichment within unique repli-ATAC-seq peaks to each doxycycline concentration and within common peaks across all doxycycline concentrations, in nascent and mature chromatin. Same at Figure 5e with motif names labeled. Each motif found is plotted by the -log10 p-value and motif family determined by HOMER. **(d)** Left: Gene expression (asynchronous, from Figure 4) of nearest genes to PAX3::FOXO1-independent (n = 687) or -dependent (n = 4415) accessible sites by doxycycline concentration in nascent chromatin. One-way ANOVA with Tukey post-hoc test was used to compare across doxycycline concentrations, ** indicates p ≤ 0.01, *** p ≤ 0.001, **** p ≤ 0.0001. All boxplots indicate median, first quartile, and third quartile and whiskers indicate 1.5x the inter-quartile range. Right: Enriched gene sets (from MSigDB Hallmark) within PAX3::FOXO1-independent or -dependent genes.

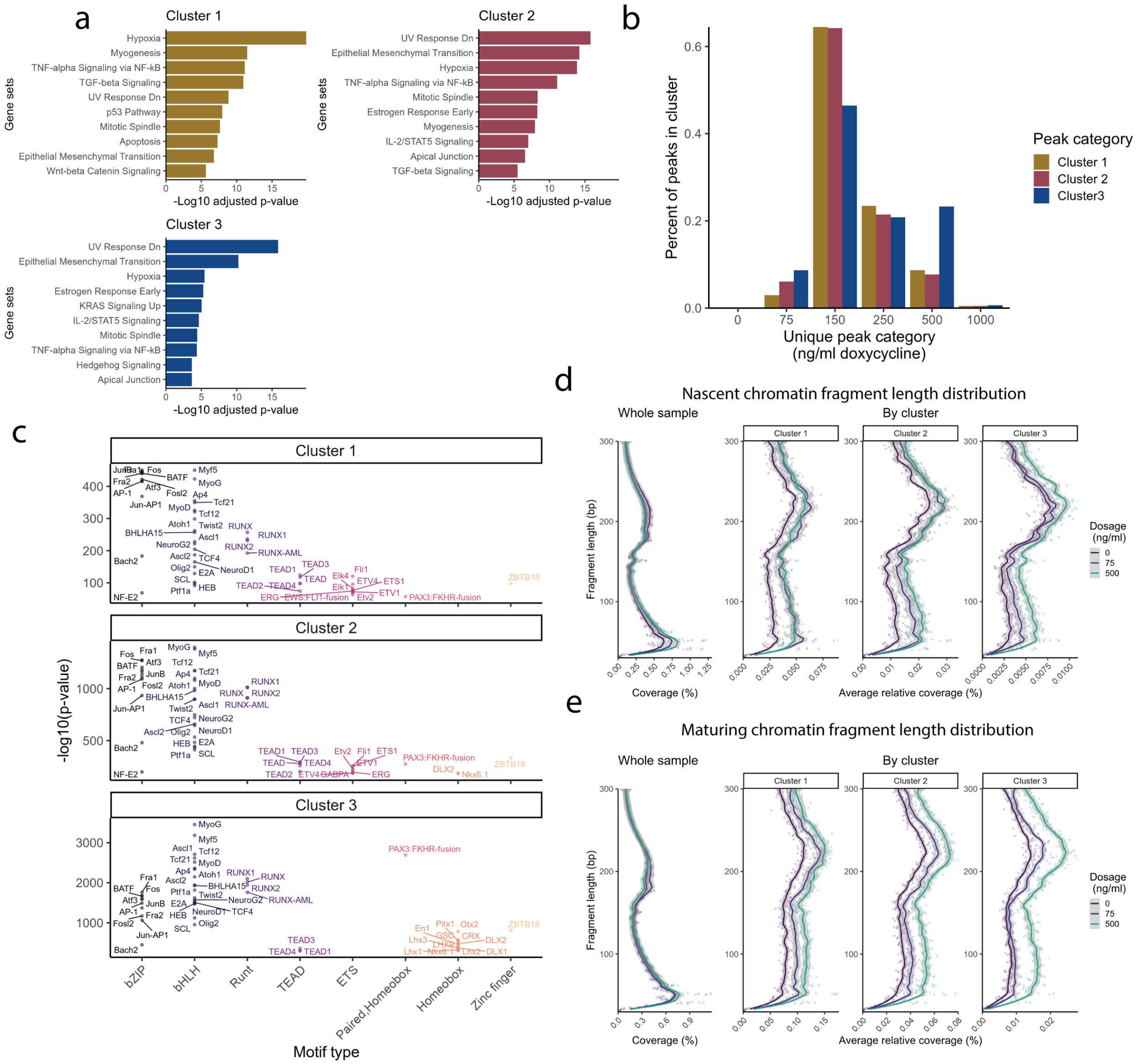


**Extended Data Figure 5. (a)** Enriched gene sets (from MSigDB Hallmark) within each cluster of PAX3::FOXO1 sites identified in Figure 6a. **(b)** Overlap of PAX3::FOXO1 sites in clusters identified in Figure 6a with sites unique to each doxycycline dosage (from Figure 3). **(c)** Motif enrichment at PAX3::FOXO1 sites in each cluster. Same as Figure 6c with motif names labeled. Each motif found is plotted by the -log10 p-value and motif family determined by HOMER. Boxplots indicate median, first quartile, and third quartile and whiskers indicate 1.5x the inter-quartile range. Nascent **(d)** and maturing chromatin **(e)** fragment length distribution from repliATAC-seq across clusters from Figure 6 and whole samples, determined using plot2DO output.

**Supplementary Data:**

| *Dosage (ng/ml)* | *Motif class* | | | | | | | |
|---|---|---|---|---|---|---|---|---|
| | **bZIP** | **Runt** | **bHLH** | **Paired,Homeobox** | **Homeobox** | **TEAD** | **Zf** | **ETS** |
| **75** | JunB, Atf3, Fosl2, Bach2, NFE2L2, Bach1 | RUNX1, RUNX2 | MyoG, Myf5, Tcf12, TCF4, Twist2 | PAX3::FOXO1 | DLX2, DLX1 | TEAD4, TEAD3, TEAD2, TEAD1 | - | - |
| **150** | Fos, JunB, Atf3, Fosl2, Bach2 | RUNX1, RUNX2 | MyoG, Myf5, Tcf12, Twist2, TCF4 | PAX3::FOXO1 | - | TEAD3, TEAD1, TEAD4 | ZBTB18, CTCF | ETS1, ETV4, ETV1, GABPA, Elk1 |
| **250** | Fos, Atf3, JunB, Fosl2, Bach2 | RUNX1, RUNX2 | MyoG, Myf5, Tcf12, TCF4 | PAX3::FOXO1 | Pitx1 | TEAD3, TEAD4, TEAD1 | ZBTB18, CTCF | ETS1, Elk1, Elk4, Etv2 |
| **500** | Fos, Atf3, JunB | RUNX1, RUNX2 | MyoG, Myf5, Tcf12, Twist2, TCF4 | PAX3::FOXO1 | Pitx1, DLX2, DLX1, Lhx2, Dlx3, Six2 | - | ZBTB18 | - |
| **common** | Fos, Atf3, JunB, Fosl2, Bach2 | RUNX1, RUNX2 | MyoG, Myf5, Tcf12, Twist2, TCF4 | PAX3::FOXO1 | DLX2, DLX1, Pitx1, Lhx2 | TEAD4, TEAD1, TEAD3 | ZBTB18 | - |

Note that there is no NR motif class column because the relevant factor is not expressed.

**Table S1.** Expressed transcription factors with motifs in dosage-specific peaksets.

**Supplemental Table 1.** Expressed transcription factors with motifs in dosage-specific peak sets. Expression determined from bulk RNA-seq data (from Figure 4).

**Supplemental Table 2.** List of genes near PAX3::FOXO1-dependent and -independent accessible sites in nascent and maturing chromatin, table_S2.xlsx
